# A scale-invariant perturbative approach to study information communication in dynamic brain networks

**DOI:** 10.1101/2021.11.24.469896

**Authors:** Varun Madan Mohan, Arpan Banerjee

## Abstract

How communication among neuronal ensembles shapes functional brain dynamics at the large scale is a question of fundamental importance to Neuroscience. To date, researchers have primarily relied on two alternative ways to address this issue 1) *in-silico* neurodynamical modelling of functional brain dynamics by choosing biophysically inspired non-linear systems, interacting via a connection topology driven by empirical data; and 2) identifying topological measures to quantify network structure and studying them in tandem with functional metrics of interest, e.g. co-variation of time series in brain regions from fast (EEG/ MEG) and slow (fMRI) timescales. While the modelling approaches are limited in scope to only scales of the nervous system for which dynamical models are well defined, the latter approach does not take into account how the network architecture and intrinsic regional node dynamics contribute together to inter-regional communication in the brain. Thus, developing a generalized scale-invariant measure of interaction between network topology and constituent regional dynamics can potentially resolve how transmission of perturbations in brain networks alter function e.g. by neuropathologies, or the intervention strategies designed to mitigate them. In this work, we introduce a recently developed theoretical perturbative framework in network science into a neuroscientific framework, to conceptualize the interaction of regional dynamics and network architecture in a quantifiable manner. This framework further provides insights into the information communication contributions of putative regions and sub-networks in the brain, irrespective of the observational scale of the phenomenon (firing rates to BOLD fMRI time series). The proposed approach can directly quantify network-dynamical interactions without reliance on a specific class of models or response characteristics: linear/nonlinear. By simply gauging the asymmetries in responses to perturbations, we obtain insights into the significance of regions in communication and their influence over the rest of the network. Moreover, coupling perturbations with functional lesions can also answer which regions contribute the most to information spread: a quantity termed *Flow*. The simplicity of the proposed technique allows translation to an experimental setting where the response asymmetries and flow can inversely act as a window into the dynamics of regions. For proof-of-concept, we apply the perturbative approach on *in-silico* data generated for human resting state network dynamics, using different established dynamical models that mimic empirical observations. We also apply the perturbation approach at the level of large scale Resting State Networks (RSNs) to gauge the range of network-dynamical interactions in mediating information flow across brain regions.

## 1 Introduction

The brain is a complex biological network in which healthy function strongly relies on the efficient transmission of information between functional entities at multiple scales [Goñi et al., 2014, Avena-Koenigsberger et al., 2018, Battaglia et al., 2011]. At the whole-brain scale, it is generally assumed that information *flows* along channels of white matter tracts connecting grey matter regions, subsequently leading to the observed brain dynamics. However, precise mapping of the relation between the structural connectome (SC), and the functional connectivity (FC) computed from empirical observations is an enduring problem in neuroscience [Suárez et al., 2020, Honey et al., 2010]. The crucial link to this mapping lies in the particulars of dynamics that evolve over the SC [Forrester et al., 2020]. The dynamical complexity of information processing units of the brain is well showcased by the plethora of dynamical models at multiple spatial scales, from single neurons [Hodgkin and Huxley, 1952, FitzHugh, 1961], to local populations [Wilson and Cowan, 1972, Stefanescu and Jirsa, 2008] and finally at the level of macroscopic brain regions [Deco et al., 2013, Naskar et al., 2021]. At the whole-brain level, neurodynamical models which evolve on the SC have been successful in providing this mechanistic link from structure to function and have been shown to be able to predict FC in varying degrees and scales [Messé et al., 2015b, Vattikonda et al., 2016, Naskar et al., 2021]. While these mathematically diverse models have been successful in capturing features such as synchronization[Cabral et al., 2011], transient dynamics [Hansen et al., 2015], neural patterns [Galán, 2008] etc, they are also limited in their generalizability to multiple spatio-temporal scales because of the lack of a conceptual framework to link how the brain dynamics of physiologically localized (or extended) tissues interact with the network ^1^. Recent advances in studies of real-world networks [Harush and Barzel, 2017, Hens et al., 2019] have proposed a theoretical framework on how to investigate the interaction of network and nodal dynamics, which can have major implications on functional capabilties of brain regions, inter-regional communication, and structure-function mapping in general.

Network-dynamical interactions have previously been implicated in the rise of emergent phenomena that transcend understanding from local dynamics, such as synchronization in oscillator networks [Strogatz, 2000], amplification in linear recurrent networks [Dayan and Abbott, 2001], occurrence of critical states exhibiting vast dynamical repertoire [Hansen et al., 2015, Sporns, 2014], as well as non-trivial signal propagation scenarios [Hens et al., 2019]. Structural brain networks have been shown to display effective connectivity asymmetries even in the resting state [Seguin et al., 2019]. Asymmetries and heterogenities in functional properties (as opposed to structural graph theoretic properties) have also been reported by studies from multiple domains [Moon et al., 2015, Gu et al., 2015, Misic et al., 2018, Seguin et al., 2019, Novelli et al., 2020]. From these studies, it is obvious that the relation of these asymmetries to the SC is seldom straightforward, and depends on the kind of dynamics that evolves over the SC, indicating a contribution of network-dynamical interactions. The introduction of dynamics on the network has also been shown to render static-network derived graph theoretic measures to be poor predictors of functional importance[van Elteren et al., 2019]. The importance of a node in a network is traditionally quantified by global and local graph theoretic measures [Newman, 2018]. The distributions and possible functional roles of these graph theoretic measures have been explored in structural and functional connectivities of both healthy and diseased brains [Rubinov and Sporns, 2010, Fornito et al., 2015, Bassett and Sporns, 2017]. For a system that relies extensively on efficient communication like the brain, identifying the regions which are central to the overall communication process is of great importance from theoretical, experimental, and clinical standpoints. Thus, a region’s contribution towards modulating arbitrary information transfer processes defines its role in maintaining overall inter-regional communication. In addition to healthy information relay, these nodes will also naturally play a crucial role in facilitating pathological spread and maladaptive responses such as diaschisis of indirectly connected regions [Carrera and Tononi, 2014, Fornito et al., 2015]. However, the inherent contribution of dynamics on the information transfer process deems purely structural centrality measures inadequate for gathering such insights into communication.

We can study the fallouts of interactions between the dynamics and the underlying anatomical network by tracing the effects of changes to a node’s activity (akin to the information) on the rest of the active network - a perturbative approach. Perturbative methods have been previously used *in silico* to gather insights into diverse aspects of brain function [Gollo et al., 2017, Deco et al., 2018, Sanz Perl et al., 2021]. The transfer of information can be captured as baseline activity perturbation that originates in one region and results in a chain of perturbations in subsequent neighbourhoods of intermediary nodes till it reaches the target (Fig.1B). This is similar to a broadcasting communication strategy [Avena-Koenigsberger et al., 2018] which has been shown to come second only to the FC in terms of behavioural predictive utility [Seguin et al., 2020]. Here, we choose to recast the information flow formalism introduced recently in network science [Harush and Barzel, 2017], to unearth interesting asymmetries in the global network influencing capabilities of distinct brain regions. Furthermore, an extension of this formalism leads to the definition of a dynamics-dependent communication centrality measure, defined as *Flow*. This measure quantifies the overall contribution of a region to information transfer events in the network. Readers should note that this method should not be confused with the Granger-Geweke causality - a proxy measure for information flow, often applied on neural time series [Dhamala et al., 2018]. The prime advantage of the perturbative approach is that the method does not alter the structure of the underlying anatomical network. This preserves structural network properties and helps in singling out dynamical consequences to the perturbations. Secondly, the close relation of this method to gauging information flow, helps in arriving at insightful metrics that quantify a region’s capability to act as an effective “mediator” of communication in an active brain network. This metric as we will discuss, can hold considerable insights into possible communication patterns supported by neurodynamical models. Lastly, the simplicity and mathematical basis of the formalism permits its easy application onto any experimental framework using perturbations, such as brain stimulation techniques. Most importantly, since the measures developed are at the level of time series obtained at any voxel or sensor, the approach is suitable for studies of single neurons to macroscopic brain recordings and is essentially scale invariant (Fig.1A).

**Figure 1.**
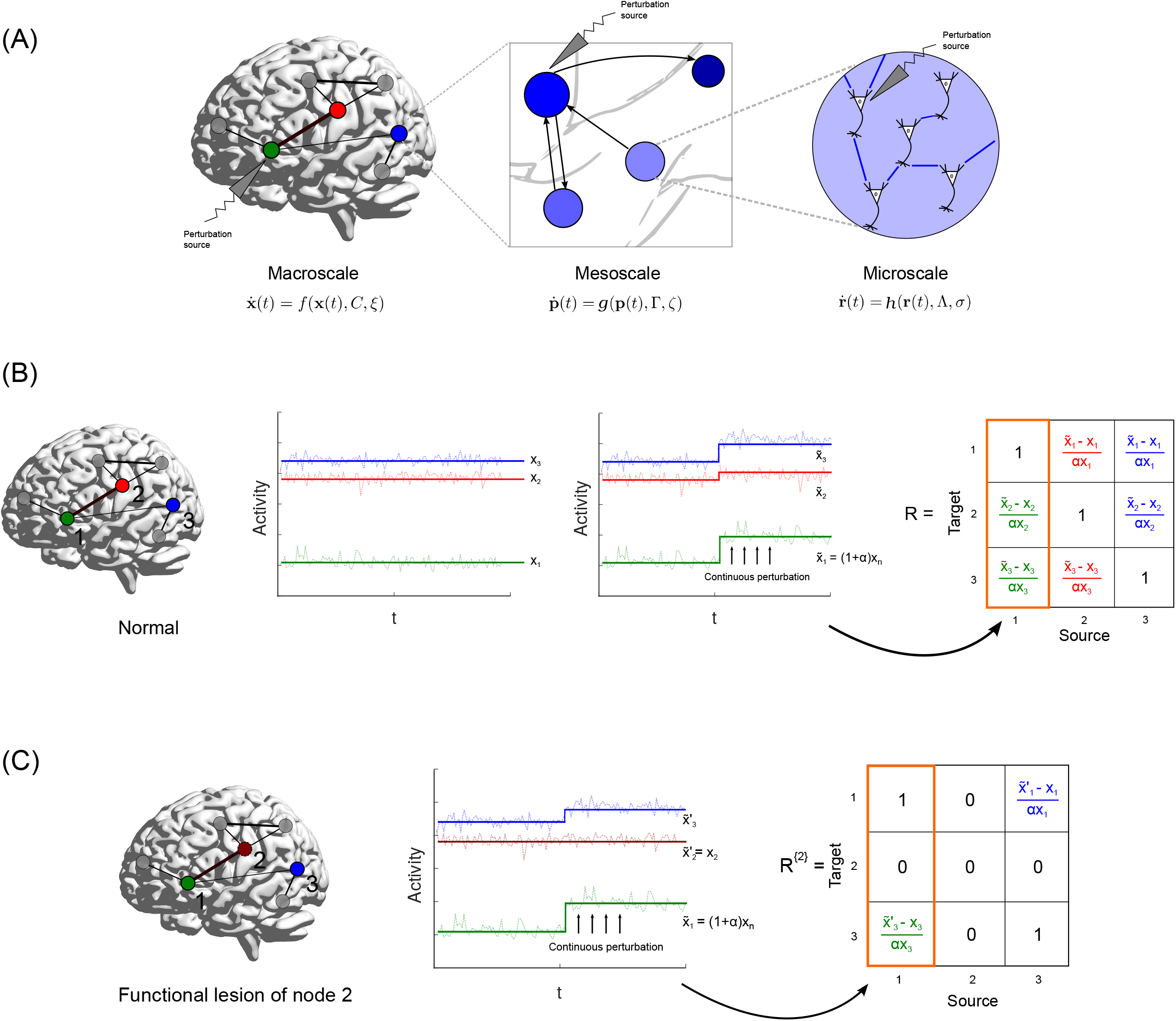
Schematic of perturbation protocol and response matrix before and after freezing node activity. **(A)** The multiscale scope of the perturbative formalism. Each scale has its own dynamical model evolving over associated networks *C, Γ*, and Λ. Perturbing the functional units results in changes in activity across the rest of the connected network, in accordance with the dynamics in that scale. This is then translated to a region’s influence and role in communication. **(B)** Nodes 1(Green), 2(Red), and 3 (Blue) are active in a whole-brain network. The steady state values at which they stabilize are given by *x_1_, x_2_*, and *x_3_* respectively. A continuous perturbation of node 1, by an amount *α* results in the activities of node 2 and 3 to stabilize at new steady state values, 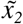 and 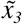 respectively. This is used to populate the first column of the linear response matrix (highlighted in orange). The colours of the entries of the response matrix *R*, corresponds to the response calculated for a perturbation of the node associated with the colour. **(C)** In order to gauge the contribution of node 2 in eliciting the responses of the other nodes, we perform a functional lesion (freeze the activity of node 2 at its original steady state). The associated columns and rows of node two are thus 0, indicating that node 2 does not respond to any perturbations. Note that this lesion of 2 leads to a different response of 3 to the perturbation of 1, with a new steady state at 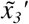.

In order to demonstrate the formalism, we resort to an *in silico* approach at the whole brain scale, using an empirically derived SC that supports established whole-brain node dynamics. This is done merely as a proof of concept and the formalism is not restricted to such approaches alone. The workflow is as follows: First, we apply the perturbation protocol on the Mean Field Model (MFM) [Deco et al., 2013], and look at steady state perturbation responses and the asymmetry that arises at the level of single brain regions (Fig.1B). We then implement a functional lesion approach to determine the contribution of single regions on the flow of information (Fig.1C). We also showcase the qualitatively different flow structures (flow-strength relationship) that arise from MFM dynamics, which we report is also preserved over two largely different datasets. We then implement the flow computation at the level of large scale neurocognitive resting state networks (RSNs), to gauge the capabilities of RSNs to mediate information flow in the rest of the brain. In order to showcase the general applicability of the method to any neurodynamical model, we also carry out the perturbation protocol on the Linear Stochastic Model (LSM) [Galán, 2008], and report associated flow patterns. We introduce the concepts of response, flow, and the associated perturbation protocols in the Methods section. Results for the *in silico* MFM over the Cam-CAN dataset is presented in the Results section, and those from implementing the protocol on an alternate dataset and dynamical model is shown in the Supplementary figures. In the final section of this work, we discuss the possible origins, implications and limitations of our findings through insights from literature and artificial networks.

## 2 Methods

### 2.1 Net influence and flow

We introduce two concepts 1) net influence of a brain sub-network and 2) the notion of “Flow”, that help illuminate and quantify network-dynamic interactions. We implement the method developed by Harush and Barzel [Harush and Barzel, 2017] on whole brain neurodynamical models whose topological architecture are constrained by empirically derived structural brain networks. In an active communicating network, information or activity between two non-neighbouring nodes is relayed by intermediary nodes. The original activity or a perturbation of the source node elicits a perturbation in the activity of its immediate neighbours, which then elicit responses in their neighbours and so on, till the effect of the source perturbation finally reaches the target. This perturbation transmission can be easily traced *in silico*, and the final effect of perturbing the source can be quantified by computing the change in activity of the target per unit change of source activity (Fig.1B).

In general, any neurodynamical model can be represented as

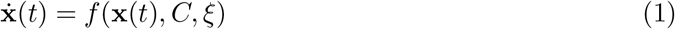

where **x**(*t*) is the vector of time-dependent activities of every region, *C* represents the network on which the dynamics evolves, and *ξ* is the noise function. If the system is stable, permitting it to evolve for long enough without any external perturbation drives it to its steady state. We can introduce a continuous perturbation held at a constant strength to a region *n*, which has the effect of driving the system to a new steady state. The steady state *linear* response matrix of a region *m* upon a small perturbation of a region *n*, such that the perturbed state of the source, 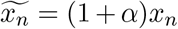 (where *x_n_* denotes the steady state activity of region *n*), is computed as

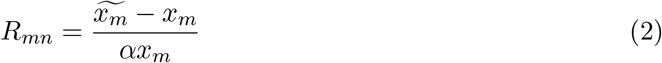

where *x_m_* is the steady state of node *m* in the unperturbed system, 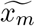 is the new steady state of *m* after perturbation, and *α* quantifies the amount of perturbation. We implement this computationally by solving the equations of all regions *m*, while maintaining the activity of region *n* at (1 + *α*)*x_n_*. Computing all pairwise responses in this manner generates the linear response matrix ^2^, *R*. The total response elicited by a region *n* is then given by

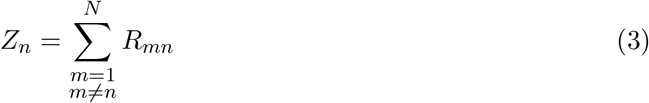

The ***net influence*** of a putative brain region can be defined as, 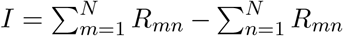. Thus, net influence essentially captures the response assymetry of a node by taking the difference between magnitude of response elicited *on* the rest of the network (sum over rows) and magnitude of response effected *by* the rest of the network (sum over columns).

Probing the effect of a region *i* on the response of *m* elicited by the perturbation of *n* requires carrying out the procedure mentioned above, but now with the activity of *i* fixed at its unperturbed steady state value *x_i_* (Fig.1C): an effective “functional lesion”. This is tantamount to *i* remaining at its stable baseline, not being perturbed by n, and thereby removing its contribution to 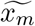 and subsequently, computing the response of all possible target source pairs by freezing the activity of *i* generates *R*^{*i*}^. However, computing *R*^{*i*}^ for all *i* is a computationally intensive process, and so we implement small perturbation approximation which only requires *R* [Harush and Barzel, 2017]:

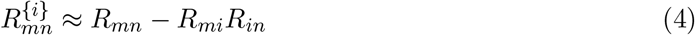

The difference between *R*^{*i*}^ and *R* is captured by the measure ***Flow*** of region *i*, *F^i^*, computed as follows:

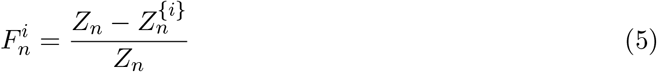

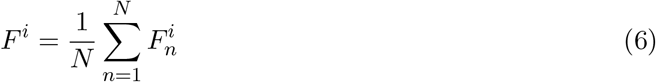

where, 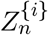 is the total response elicited by perturbing *n*, when the activity of *i* is frozen.

To demonstrate the insights gained from the two concepts, ***Net influence*** and ***flow***, we extracted structural brain networks from publicly available datasets that incorporated diffusion weighted imaging (DWI) and functional magentic resonance imaging (fMRI) modalities. Dynamical models of neural ensemble activity at macroscopic scales - Mean field model (MFM) and the linear stochastic model (LSM), can be used to simulate the node dynamics of brain networks (discussed in detail in section 2.3). Using an optimatzation of approach of functional conncetivity distance [Vattikonda et al., 2016, Naskar et al., 2021], we tune the network parameters to be set in a regime to capture the empirical resting state functional connectivity (rs-FC) to maximum extent. Thereafter, using the simulated time series we compute the net influence and flow for all brain nodes (parcels) with respect to the whole brain, and individually for each resting state brain network.

### 2.2 Datasets

The generality of the two measures - *net influence* and *flow*, were validated on two seperate human neuroimaging datasets obtained from Cambridge Centre for Ageing and Neuroscience (Cam-CAN) cohort [Shafto et al., 2014, Taylor et al., 2017] and the Nathan Kline Institute (NKI)/Rockland sample (available in the UCLA Multimodal Connectivity Database (UMCD)) [Brown et al., 2012]. Different parcellation schemes were used on each of the data sets to further showcase that the outcomes do not depend on the extent of brain parcellations that govern the size of the network analyzed.

#### 2.2.1 Cam-CAN

Empirical Diffusion Weighted Imaging (DWI) data of 40 healthy participants (19 males; 21 females, total age range 18-38), sampled from the Cambridge Centre for Ageing and Neuroscience (Cam-CAN) cohort[Shafto et al., 2014, Taylor et al., 2017] chosen for an earlier work [Naskar et al., 2021] was used to construct the structural connectivity (SC) matrix. Cortical gray matter was parcellated into 150 ROIs using the Destrieux parcellation [Destrieux et al., 2010] and the subject-wise SC matrix was generated using an automated pipeline by [Schirner et al., 2015], and was averaged over the 40 subjects to generate the group-averaged SC matrix. A curious reader may refer to the original article for additional details about the preprocessing steps used [Naskar et al., 2021].

The resting state functional connectivity (rs-FC) matrix was required for determination of model parameters that best fit empirical data. We use a 150 region Destrieux parcellated rs-FC matrix that was also used in the earlier article [Naskar et al., 2021]. The rs-FC was generated by computing the pairwise Pearson coefficient between all pairs of z-transformed BOLD timeseries, for 40 subjects, and then averaged to generate the rs-FC used for empirical model validation.

#### 2.2.2 NKI

Diffusion Tensor Imaging (DTI) based structural connectivity matrices and rs-FC/Pearson correlation matrix of 171 healthy participants were obtained from the Nathan Kline Institute (NKI)/Rockland sample as made available in the UCLA Multimodal Connectivity Database (UMCD) [Brown et al., 2012]. The participants consisted of 99 males and 72 females, with a total age range of 4-85. We used the 171 participant group averaged SC and rs-FC for the analysis. Cortical gray matter was parcellated using the Craddock 200 atlas [Craddock et al., 2012]. Elements of the SC represented the number of white matter tracts between gray matter parcels. The reader may also refer to the original work [Brown et al., 2012] for details regarding structural and functional connectome preprocessing.

Prior to running the models on the NKI SC, the SC was scaled down by dividing every element by the maximum value found in the original SC consisting of white matter tract numbers between regions. This is only done to bring the critical range of the MFM free parameter to a value that is comparable to that found for the Cam-CAN SC, and does not bring about any topological changes to the SC.

### 2.3 Neurodynamical models

A number of neurodynamical models have been formulated to simulate whole brain resting state activity from DTI/ DWI based structural connectome. Some of the notable models include the Vector Auto Regressive (VAR) model [Tononi et al., 1994, Messé et al., 2014, Messé et al., 2015a], Linear stochastic Model [Galán, 2008, Goñi et al., 2014, Hansen et al., 2015], Wilson-Cowan oscillator [Wilson and Cowan, 1972], Kuramoto oscillator [Cabral et al., 2011, Abeysuriya et al., 2018],and Mean Field Model [Deco et al., 2013, Hansen et al., 2015, Vattikonda et al., 2016, Naskar et al., 2021] among others. In order to depict the variety of network-dynamical interactions supported by an underlying SC, we use two models that we believe are far apart in terms of their dynamical complexities: the Linear stochastic model and the Mean Field Model.

All stochastic neurodynamical models were numerically integrated using the Euler integration algorithm [Mannella, 2002], with a timestep of 1ms. Simulations to arrive at empirical best fit model parameters were carried out for 315.2s. The first step of the noiseless perturbation protocols were run for 60s, with a time step of 1ms. This was done to ensure that the system stabilised at its steady state. The second step, the perturbation, was carried out for 5s, with a time step of 1ms. We ensured that the simulation times were long enough for steady state stabilization. The intial value of the dynamical variable for all the models were sampled from a uniform distribution bounded between [0, 1].

#### 2.3.1 Mean Field Model

The dynamic Mean Field Model (MFM) is obtained by carrying out a mean field reduction [Deco et al., 2013] of the spiking neuron model [Deco and Jirsa, 2012]. The MFM has also been shown to replicate features of empirical functional state transition dynamics, in addition to being able to predict time averaged FC [Hansen et al., 2015]. The dynamics of the MFM nodes are given by:

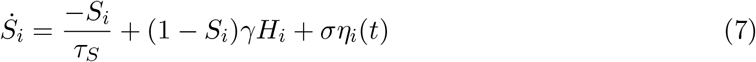

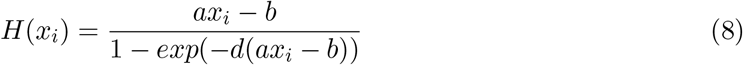

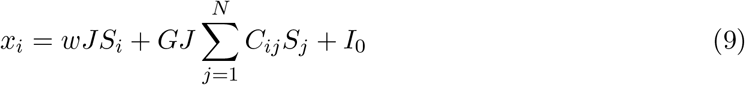

where *S_i_* is the NMDA synaptic gating variable of the *i^th^* node, *H*(*x_i_*) is the firing rate function for the input *x_i_*, to the node *i*. Following [Deco et al., 2013], the fixed parameters are the local excitatory recurrence *w* = 0.9, synaptic coupling *J* = 0.2609(nA), overall external input *I*_0_ = 0.3(nA), kinetic parameters *γ* = 0.641/1000 and *τ_S_* = 100(ms), and firing rate function parameters *a* = 270(n/C), *b* = 108(Hz), and *d* = 0.154(s). *σ* = 0.001 is the noise amplitude and *η_i_*(*t*) is random number sampled from a normal distribution.

In contrast to the LSM, nodes following MFM dynamics can exhibit bistable and non-symmetric steady states. The rich dynamical profile of the MFM can be explored by varying the global scaling parameter, G, which is a free parameter in the model. While the low-activity “spontaneous fluctuations” state is highly stable for lower values of G, increasing it pushes the system to a bistable regime. Further increase of G renders the spontaneous state unstable.

#### 2.3.2 Linear Stochastic Model

The Linear Stochastic Model (LSM)[Galán, 2008] is a simple neurodynamical model that can be obtained by removing the saturation function and inhibitory population of the Wilson-Cowan model[Wilson and Cowan, 1972]. The dynamics follow:

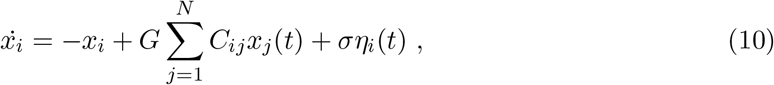

where *x_i_*(*t*) is the activity of region *i*, *G* is the global scaling parameter, *C_ij_* is an element of the structural connectivity matrix, *σ* is the noise amplitude, and *η_i_*(*t*) is a random number from a Gaussian distribution with zero mean and unit variance. All nodes following LSM dynamics have two possible steady states, depending on the value of *G*: zero for 0 ≤ *G* < *G_crit_*, and +∞ for *G* > *G_crit_*. In order to keep noise from pushing the system into the divergent regime, we first determine *G_crit_* by carrying out a noiseless simulation by varying *G* in steps of 0.01, for 20 trials. Once *G_crit_* is determined, we set the noise amplitude as *σ* = *G_crit_* − *G* [Hansen et al., 2015] for all successive stochastic runs of the model.

### 2.4 Synthesis of BOLD signals and parameter space identification

Synthesised neural activity from the dynamical models were used to generate BOLD time series using a Balloon-Windkessel hemodynamic filter [Friston et al., 2000, Friston et al., 2003, Cabral et al., 2011]. The parameters for the model were set as per [Friston et al., 2003]. The data of the first 39400ms were discarded to remove transients. Generated BOLD signals were then downsampled at 1.97s to match the sampling time of the empirical BOLD signals. The synthesised BOLD signals were then z-scored and the pairwise Pearson correlation was computed, to generate the simulated rs-FC.

Parameter space for the model for each data set can be tuned using an approach developed by earlier studies [Vattikonda et al., 2016, Naskar et al., 2021] where the difference between the model predicted FC and the empirical FC was quantified using the FC distance measure, given by:

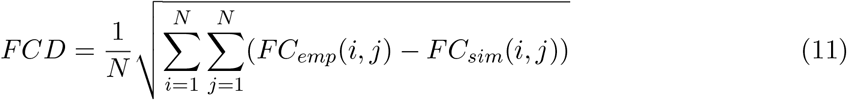

The value of *G* at which minimum FCD is observed is refered as the *G*\*.

### 2.5 Computing Net *influence* and *Flow*

Net influence, *I*, and flow, *F*, were computed for each node of the entire brain from the simulated resting-state time series. For the Cam-CAN data, the number of nodes were 150 based on the Destrieux parcellation scheme. For the NKI data, number of nodes were 188 (presented only in Supplementary Information).

Node dynamics was simulated for both model (MFM and LSM) progressing over a diffusion weighted imaging (DWI) based structural substrate as follows: First, the model is initialised randomly and allowed to noiselessly evolve over a period of 60 seconds (to ensure that the steady state is reached), with a timestep of 1ms. Second, each node is chosen as a source, n, and its value is reset to (1 + *α*)*x_n_*, and the remaining nodes of the system is initialised at their respective steady state values. The system is then allowed to evolve for 5 seconds with a timestep of 1ms, and the final steady state values of the nodes 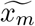 was stored.

To study how the net influence and flow behave for individual resting state sub-networks, Cam-CAN data in Desteriux parcellation was transformed to Schafer-Yeo-7-network classification [Thomas Yeo et al., 2011]. This was done by first computing the Euclidean distances between the MNI coordinates of a node in the Destrieux atlas with that of each node in the Schaefer-Yeo 7-network atlas (1000 parcel resolution). Then, the network associated with the Destrieux node is set as that of closest Schaefer-Yeo node (minimum Euclidean distance).

Once parcellated in Scahfer-Yeo scheme, each node was assigned to one of 7 large scale RSNs. The non-functionally lesioned linear response matrix is then calculated using equation 2. Just as in the case of the single node flow computation, every node in the RSN of interest is then functionally lesioned. As the total number of RSNs is small, we forego the use of the approximation (equation 4) and the perturbation-response calculation was carried out for every pairwise combination of nodes not belonging to the RSN, to yield *R^net^*. The contribution of the RSN to the overall flow from a single node is calculated using equation 5. The flow per node for the RSN is then obtained as:

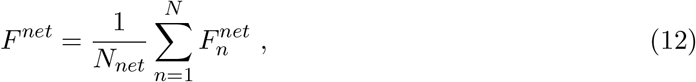

where *N_net_* is the number of nodes in the RSN.

## 3 Results

*Net influence* and *Flow* were computed from simulated resting-state time series generated from the whole-brain connectome simulations where each node can be modeled according to a neurodynamical model, e.g., Mean field model (MFM) or Linear Stochastic Model (LSM). The structural connectivity (SC) matrix for the whole-brain model was generated from the Cam-CAN database, and resting-state functional connecivity (rs-FC) from empirical data was used to optimize model parameters (see sections 2.3—2.5 for details). In this section, we first report the results of undertaking a perturbation protocol node-by-node and subsequently estimating the net influence for each node in the whole-brain connectome. Second, we computed flow, which implements the functional lesion step, together with the standard perturbation protocol, and highlights brain regions that are crucial to the overall transfer of information in the brain. The final result involves computing the flow by functionally lesioning each resting state network (RSN) as opposed to single nodes. In all three analyses, the net influence and flow are studied as a function of local neighborhood properties (node strength). Further, we inverstigated how this relationship changed when the free parameter *G* of MFM was varied, with *G*\* corresponding to the the value of *G* that best fits empirical data. Previous studies [Deco et al., 2013, Naskar et al., 2021] have shown that healthy resting state brain dynamics exhibits maximum metastability occuring at a critical value of *G*\*, around a weak global coupling regime. Following the same line of reasoning, any critical or threshold-like behavior of metrics like net influence and flow at *G*\* demonstrates sensitivity to critical changes in global dynamics of the system under ivestigation and hence qualifies as a ground truth validation. The computation of net influence and flow for other values of *G* (representative of low activity, bistable, and high activity regimes - Fig.2) further depicts various information transfer scenarios in a network that can be captured by the two measures.

**Figure 2.**
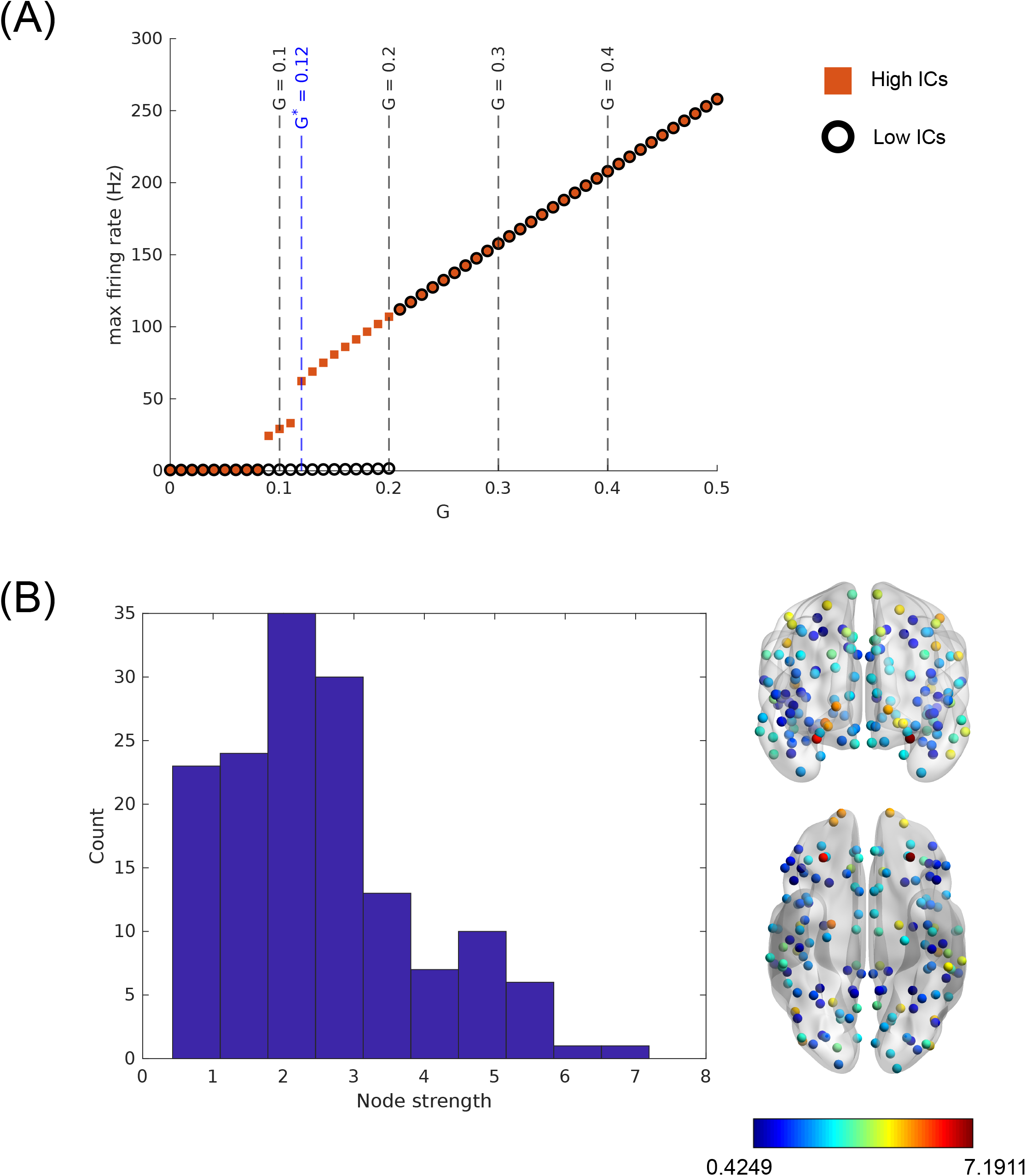
Mean Field Model (MFM) dynamical landscape for the Cam-CAN dataset. **(A)** The figure shows the dynamical landscape of the MFM as a function of the global scaling parameter *G* (orange squares = High initial conditions (ICs) (0.3 ≤ *S*(*t* = 0) ≤ 1.0), black rings = low ICs (0 ≤ *S*(*t* = 0) ≤ 0.1)), quantified as the maximum firing rate among all the nodes in the network. Figure shows results for 10 random trials at each value of *G* (*ΔG* = 0.01). The values of *G* for which the response asymmetries and flows are calculated are marked by the dashed lines. The blue dashed line corresponding to *G*^*^ = 0.13 is the value of *G* for which the model maximally conforms to empirical data. **(B)** Node strength distribution of the Cam-CAN dataset (*left*) with 150 nodes of the Destrieux atlas, and associated BrainNet plots [Xia et al., 2013] where colour maps to node strength (*right*).

While the main text presents results for the MFM evolving on the SC derived from the Cam-CAN dataset, as a demonstration of generality, the entire pipeline was repeated when the neurodynamical model in use was LSM, and the SC (and rs-FC used for parameter optimization) was estimated from NKI database. These results are presented in the Supplementary figures.

### 3.1 Perturbation unearths asymmetries in influence capabilities

Non-zero net influence following a perturbation protocol captures the response asymmetry for a particular node of the network, and is a function of node strength (sum of fibre densities from all its connections). We used the SC reported by an earlier study [Naskar et al., 2021], which can be refered to for computation of the SC, whose elements quantify the fiber densities between cortical regions. Interestingly, the relationship between the net influence and the node-strength computed on MFM simulations on the Cam-CAN SC, displays a critical dependence on *G* (Fig.3). At critical *G*\* = 0.12, overall response asymmetry is maximum. As *G* is increased, we observe that the magnitude of asymmetry decreases, with the response of regions to perturbations from the rest of the network becoming comparable to the response it elicits in the rest of the network through its own perturbation. The topological profile of the asymmetry, which is captured by its relation to the node strength shows that high strength “hubs” unequivocally exert a positive influence on the rest of the network. The influence of the nodes of low strength (the “periphery”) on the other hand, although negative, depends on *G*, with the most negative influence at *G*\* shown by intermediate strength nodes. This non-monotonic strength dependence of the net influence is observed in the bistable regime. From visual inspection it is evident that the response asymmetries are non-lateralized as depicted by the BrainNet plots [Xia et al., 2013] in Fig.3. For the NKI data set, similar dependence with *G* was observed as well (Supplementary Fig.S3).

**Figure 3.**
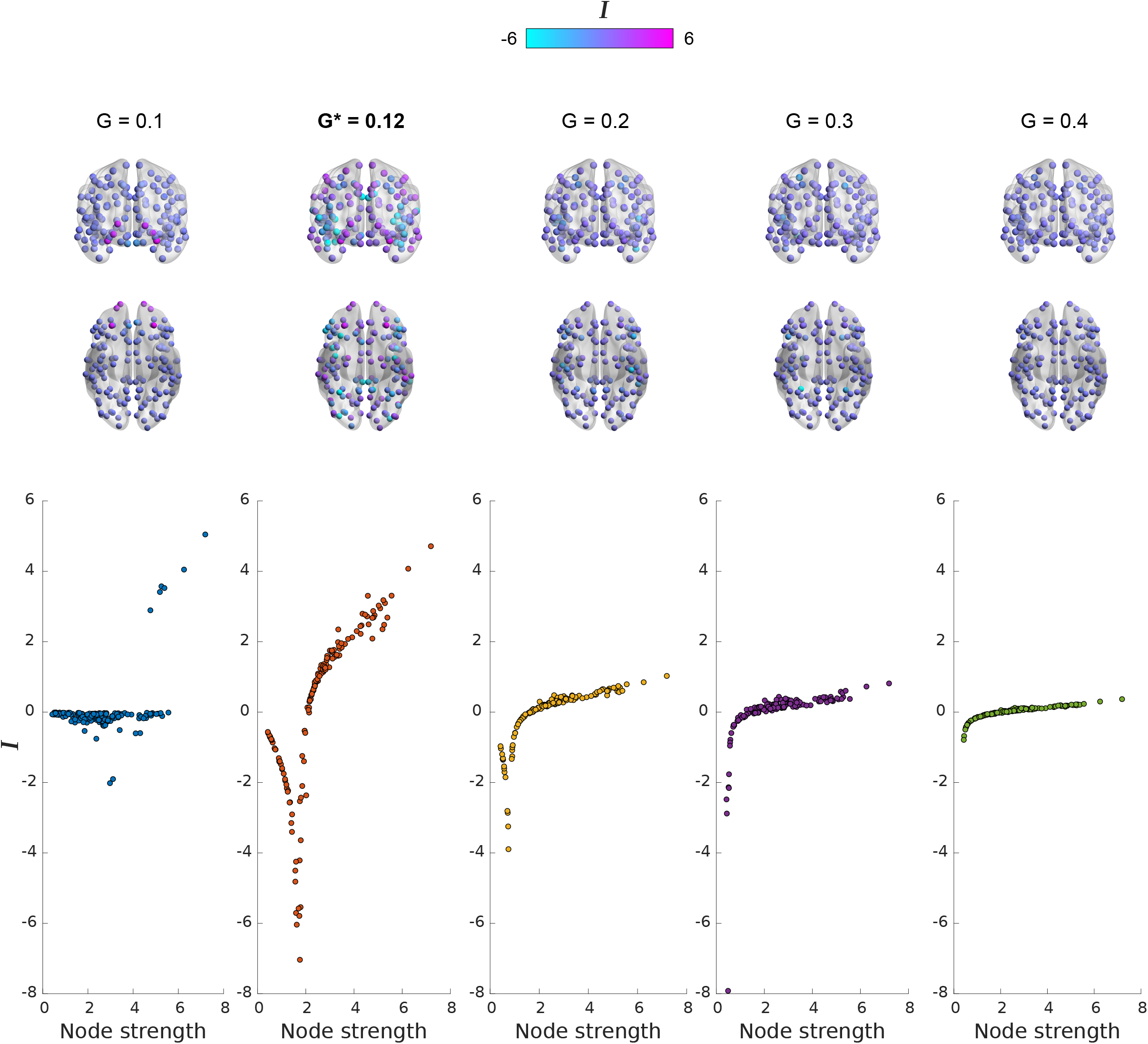
Response asymmetry. Response asymmetries, quantified by the net influence of nodes, *I* = *Σ*_m=1_*R*_mn_ - *Σ*_n=1_*R*_mn_, as a function of node strengths. The values of the global scaling parameter for which the asymmetries are computed are shown in Fig.2. Anterior and ventral view BrainNet plots depict the inter-hemispheric symmetry of net influence distribution, and the colour maps to the associated value (maximum colour value set to 6 for clarity). Overall response asymmetry is maximized at *G*^*^ = 0.12 and reduces as *G* is increased.

### 3.2 Flow is maximized for nodes of intermediate strength

The flow structure (flow-node strength relationship) computed for the MFM shows a clear dependence on the scaling parameter *G* (Fig. 4). Flow patterns varied from a hub-dominated flow structure for *G* < *G*\* to an increasingly periphery dominated flow structure for *G* > *G*\*. At the critical *G*\*, we find that flow isn’t strictly periphery dominated, and is rather maximized for nodes that have an intermediate strength. This trend is observed even at the edge of the bistable regime (*G* = 0.2), albeit for lower strength nodes, indicating a transition towards a periphery dominated flow structure with an increase in *G*.

**Figure 4.**
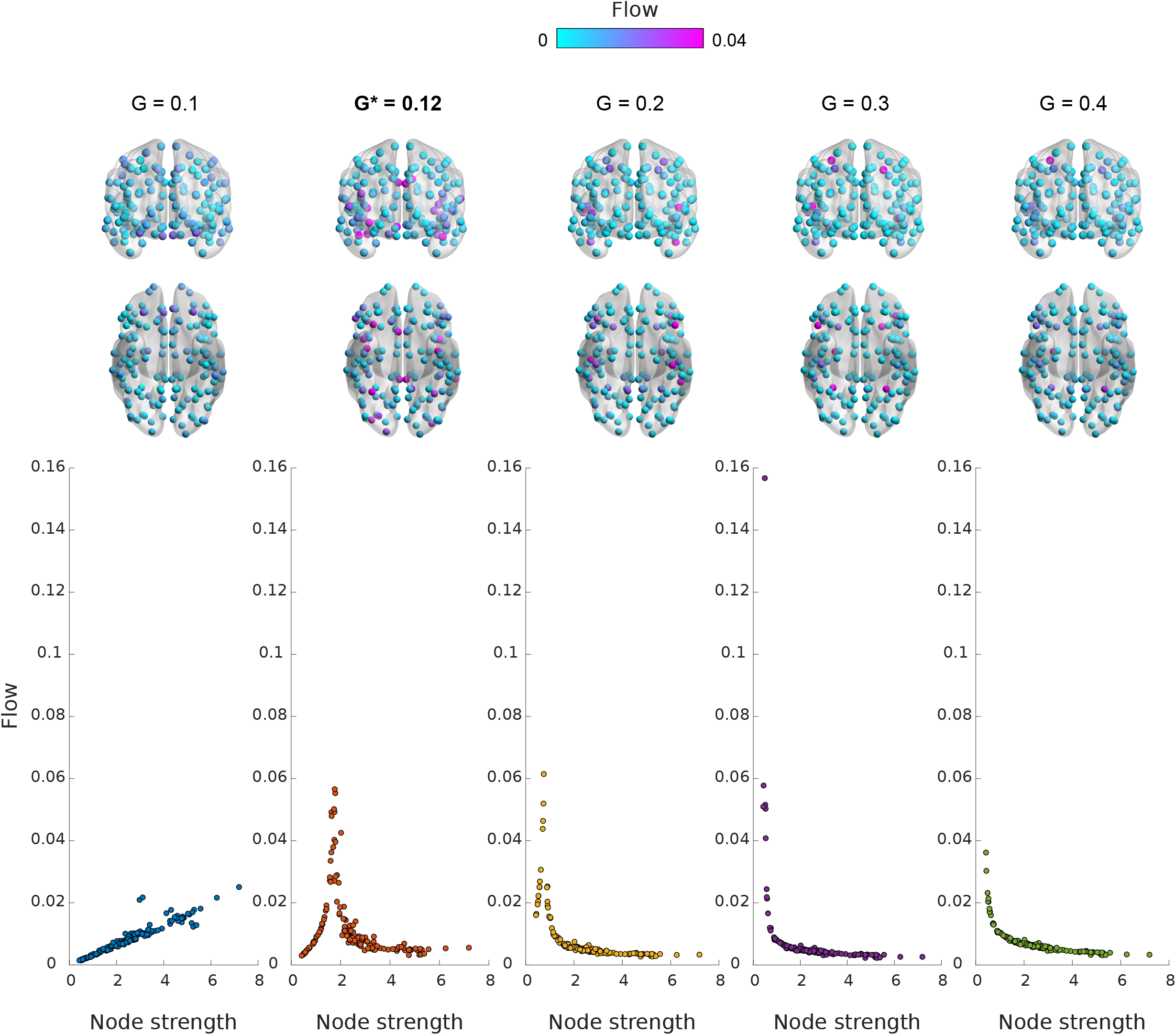
Node-level flow structure. Flow through a node, quantified as the effect of freezing the activity of node on the magnitude of responses elicited on the rest of the network, as a function of node strengths. The values of the global scaling parameter for which the flow values are computed are shown in Fig.2. Anterior and ventral view BrainNet plots depict the inter-hemispheric symmetry of flow distribution, and the colour maps to the associated value (maximum colour value set to 0.04 for clarity). Flow is maximized at intermediate strength nodes for *G*^*^ = 0.12, but moves further towards the periphery as *G* is increased.

We also observe that, for *G* ≥ *G*\* the regions displaying high flow also display a net negative perturbative influence on the network. This relation between the response asymmetry and flow structure is not observed for *G* < *G*\*. Identical pipeline run on the NKI data set (Supplementary Fig.S4) yields qualitatively similar patterns as seen in Fig.4.

### 3.3 Flow through Resting State Networks (RSNs)

We implemented the flow protocol at the level of large scale RSNs to gauge the role played by them in mediating information flow in the brain, for different values of G. The change in flow with the variation of G shows no universal trend when all RSNs are considered. However, the Salience/Ventral Attention, Frontoparietal control, Default Mode, Somatomotor, and Limbic networks show a maximization of flow at *G*\*, whereas the flow increases with *G* for the Visual network and decreases with *G* for the Dorsal Attention network (Fig.5).

**Figure 5.**
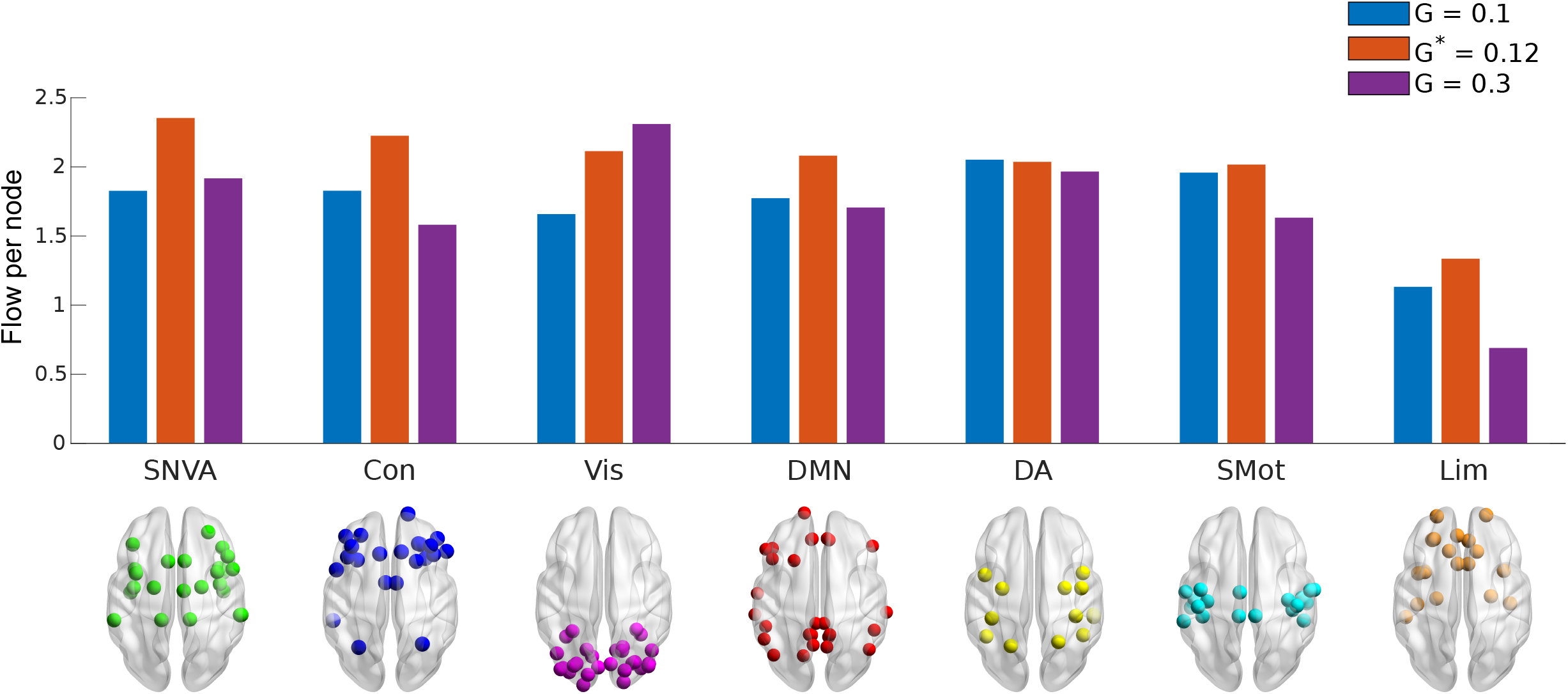
Network-level flow structure. The flow per node of a large scale Resting State Network (RSN), quantified as the effect of freezing the activity of nodes comprising an entire RSN on the magnitude of responses elicited on the rest of the network. Values are scaled by the inverse of the number of nodes in the network, to obtain the flow per node. Each coloured bar corresponds to the flow for an associated value of the global scaling parameter (*G* = 0.1, *G*^*^ = 0.12, and *G* = 0.3). The networks are arranged in descending order of flow per node at *G*^*^ for clarity. Con - Frontoparietal Control network, SNVA - Salience/Ventral Attention network, DMN - Default Mode network, Vis - Visual network, Lim - Limbic network, SMot - Somato-motor network, DA - Dorsal Attention network

At *G*\*, regions in the Salience/Ventral Attention network (in accordance with Schaefer-Yeo’s 7-network classification [Thomas Yeo et al., 2011, Schaefer et al., 2018]) have the highest contribution to the flow in the network, with the regions in the Limbic system having the lowest value of average flow per node.

## 4 Discussion

In this article, we propose a scale-invariant approach inspired by previous work in network science [Harush and Barzel, 2017], that quantifies the interaction between brain network structure and the dynamics exhibited by the constituent nodes by studying the response of the individual brain regions (nodes) as well as sub-networks (a well-defined collection of interacting nodes) established in the literature to controlled perturbations. We apply this technique on simulated data generated from an empirically derived whole-brain connectome (from the Cam-CAN cohort) where constituent nodes follow Mean Field Model (MFM) dynamics, for purposes of demonstrating its applicability. We find that the dynamically rich MFM is capable of exhibiting a wide range of qualitatively different network-dynamics interaction that can be captured in the form of response asymmetries and flow structures, as a function of the topological features of the network (node strength) and the model parameters. Importantly, the metrics introduced here, *net influence* and *flow* are sensitive to the free parameter *G*, that controls the global coupling in the network [Deco et al., 2013, Naskar et al., 2021], thus, providing scope for ground truth validation. We could replicate the results when a different cohort and parcellation scheme (NKI Rockland) was used to derive the structural connectivity (Supplementary Fig.S3, Fig.S4). Interestingly, when the neurocognitive model used was replaced with linear stochastic model (LSM) to simulate resting state brain dynamics, the non-linear dependence of net influence and flow with node strengths couldn’t be observed (Supplementary Fig.S5). This illuminates the implications of node dynamics on information flow, despite their abilities to synthesise data that conform to the empirical ground truth.

Beyond the demonstration on a specific model such as MFM, the broad objective of this study is to showcase the applicability of this technique on systems of any scale, as long as it is stable and responsive to external perturbations. This makes it a powerful tool with great potential in a multi-scale domain like neuroscience. Furthermore, the insight afforded by this formalism using just the response of regions to perturbations, opens up venues of experimental validation of communication properties of computational models and identification of flow centres through stimulation studies. Moreover, the inverse problem of what form of node dynamics can lead to observed response asymmetries and flow patterns can greatly narrow down the search for candidate models of neural dynamics. We would also like to delineate the origins of our proposed method from two popular approaches in Neuroscience that attempts to quantify information processing among brain regions, the dynamic causal modelling (DCM) [Friston et al., 2003] and Granger-Geweke causality [Dhamala et al., 2018]. These methods rely on statistical covariation of brain activities between two brain regions which can be taken as a proxy to information communication. Our proposed approach on the other hand directly quantifies information communication as a product of the network architecture and dynamical complexity of the system.

### Response and flow asymmetry along the core-periphery axis

The significance of the core-periphery axis has been pointed out by studies on various fronts, such as perturbation transfer [Gollo et al., 2017, Mišić et al., 2015], ignition dynamics [Castro et al., 2020], global communication [van den Heuvel et al., 2012, Harriger et al., 2012], task learning [Bassett et al., 2013], multi-layer frameworks [Battiston et al., 2018], and timescales of associated dynamics [Gollo et al., 2015]. Furthermore, neuropathological conditions have been widely associated with changes to the structural core [Crossley et al., 2014, Van Den Heuvel et al., 2013, Stam, 2014, Fornito et al., 2015], as opposed to the periphery. The response asymmetry reported in this work shows a clear distribution along the core-periphery axis, with peripheral “follower” nodes predominantly exerting a negative influence on the network and hub “influencer” nodes exerting a positive influence on the network (Fig.3). A net positive influence translates to the ability of a node, upon perturbation, to elicit a greater response on the rest of the network, than the response of the node itself to the perturbation of the rest of the network. Thus, any perturbation in the activity of a hub region (that can possibly result from a neuropathology, exogenous stimulation etc.) will have considerable effects on the rest of the network, whereas the rest of the network will have a considerably weaker effect on it. Our results thus agree with previous studies and suggest that hub regions make ideal high impact targets for disruption of network function.

Neurodynamical models have been shown to display critical regimes associated with the onset of dynamical richness and spatiotemporal organization of nodes into resting state networks [Deco et al., 2011, Haimovici et al., 2013, Deco and Jirsa, 2012]. Models that operate at these critical working points are also found to conform to a great degree to empirical resting state data. Our findings point out that this critical regime is also associated with the maximum overall response asymmetry. The magnitude of this asymmetry defines a clear hierarchical structure of influence of nodes, that is lost as we move away from this criticality. The functional implication of this hierarchy varies on two dimensions: the perturbative impact and role in communication. While perturbation of the influencer nodes exerts the maximum impact on the network as previously mentioned, the follower nodes are responsible for the transfer of these perturbations to different network targets. It is thus possible to limit the extent of this impact by targeting the nodes which contribute the most to this communication process.

### Response asymmetry: a direct consequence of network dynamical interactions

Although it seems intuitive to expect the response of directly connected bi-directional weight symmetric nodes upon perturbation to be equal, this study provides clear evidence to the contrary. In order to clearly illustrate the origin of this asymmetry, we design a simple toy network as depicted in Fig.6. on which we run a similar perturbation protocol using the MFM (with *G* = 1), as used in the empirically derived brain networks. In Fig.6A, the response elicited by region 1 on 2, although directly connected, is not solely a direct influence, but a product of the perturbation reaching 2 through all possible paths from 1 to 2, i.e. it is a network effect. The perturbation from 1, in addition to reaching 2 through the direct edge connecting them, also traverses through the rest of the network, via 1’s other outgoing edges. The perturbation, upon reaching a node, causes a change in its activity, and this change is then transmitted to successive neighbourhoods. Thus, the perturbation from 1 impinges upon 2 through all of 2’s incoming edges. This implies that the response of a region depends on its local neighbourhood, and not on a single path between source and target. Nodes with similar neighbourhoods should thus respond similarly to a perturbation. We see this clearly in Fig. 6B, where nodes 1 and 5, and nodes 2 and 3 have similar local neighbourhoods, as a result of which the perturbation of node 4 elicits a similar response on the pairs.

**Figure 6.**
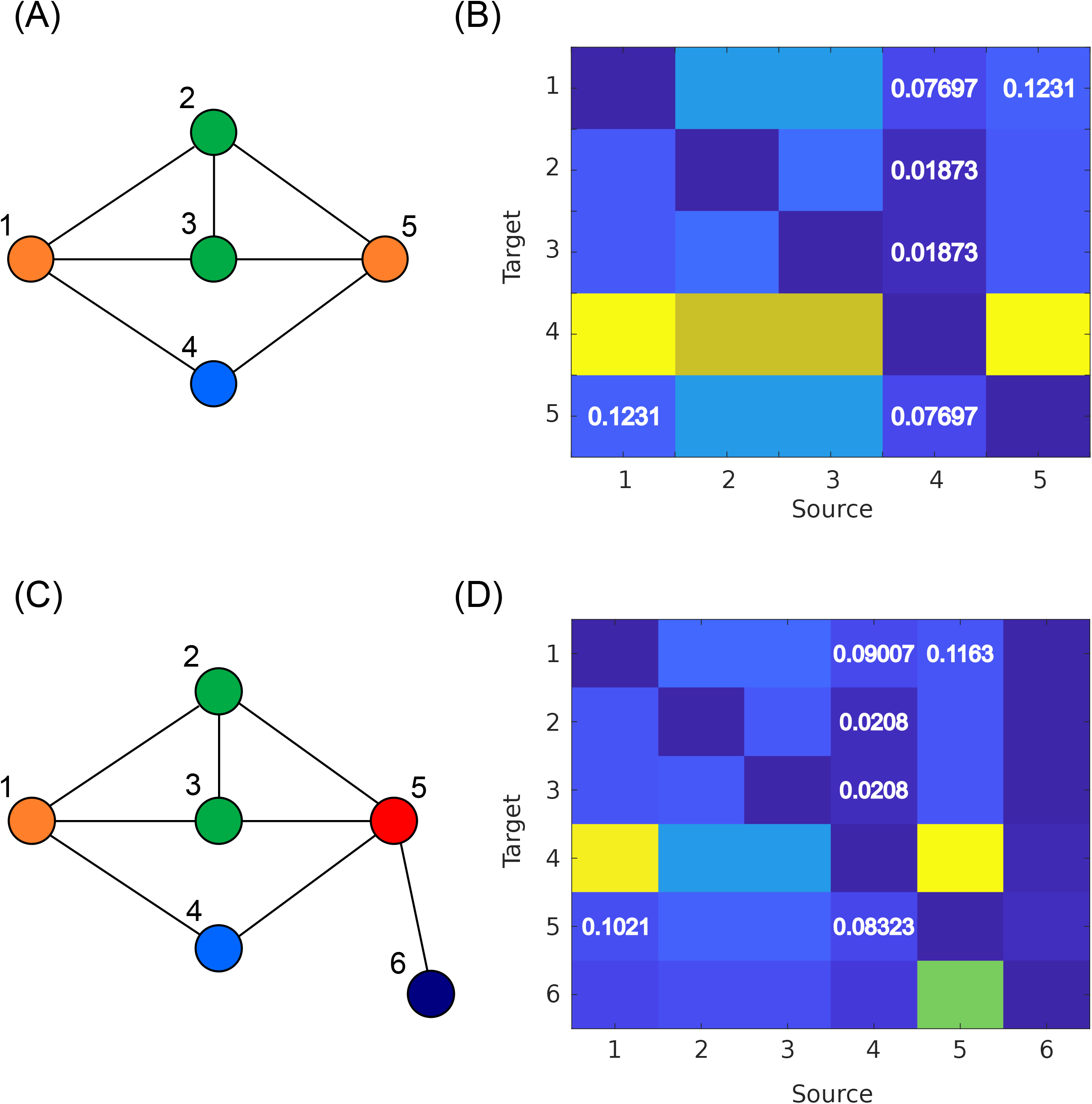
Effects of local neighbourhoods on perturbation response. **(A)** A toy network comprising 5 nodes is subjected to the perturbation protocol. The colours of the nodes are indicative of their local neighbourhoods. Nodes with the same colour have exactly similar local neighbourhoods. **(B)** Computation of the linear response matrix by successively perturbing each node shows that the effect of perturbing nodes with similar neighbourhoods have similar responses. Perturbation of node 4 elicits the same response on nodes 1 and 5, and nodes 2 and 3, which have exactly similar neighbourhoods. Moreover, the perturbation of node 1 results in a response of node 5 which is equal to the response elicited in node 1 by the perturbation of node 5 i.e. there is response symmetry. **(C)** Introduction of a sixth node in the network, changes the local neighbourhood of node 5. **(D)** This breaks the response symmetries observed in (B). The responses of nodes 2 and 3 however are preserved, as their local neighbourhoods remain unchanged. Response asymmetry thus depends only on the local neighbourhood and not on global network features.

An addition of a sixth node which is connected to node 5 (Fig.6C) breaks the similarity of local neighbourhoods of 1 and 5, and we thus see that an asymmetry arises (response of 5 upon 1’s perturbation is unequal to the response of 1 to 5’s perturbation: Fig.6D). The inter-hemispheric similarity of response asymmetry (Fig.3) and flow patterns (Fig.4) can be attributed to the similarity in local neighbourhoods of regions across hemispheres. The dynamical relation of this asymmetry borne out of local neighbourhood variability can be attributed to the form of the interaction terms in the dynamics, which for most neurodynamical models depend only on the immediate neighbourhoods of nodes. A dynamical form in which a region’s interaction with the network extends to beyond its immediate neighbourhood would imply that response asymmetry would arise with variability of not just the local neighbourhood, but the extended neighbourhood instead.

### Flow hubs are mediators of information transfer

Modelling communication or signalling dynamics in brain networks is one of the major problems in Neuroscience with far reaching implications, as communication underlies the healthy function of a functionally heterogeneous connected entity such as the brain [Avena-Koenigsberger et al., 2018, Graham et al., 2020]. Some of the popular models of communication include navigation, shortest path routing, random walk, percolation based transfer, and broadcasting. Communication models have also been shown to bolster the behavioural and functional predictive capability of structural connectomes [Seguin et al., 2020]. The neurodynamical models considered in this study are akin to the broadcasting communication strategy, which is a generalized parallel communication paradigm that assumes information flow to occur through all possible paths from every node - which we visualise as successive perturbations of neighbourhoods. The broadcasting paradigm has the added advantage of not requiring constituent nodes to have any prior information regarding global topology, and signal transfer solely occurs through the interaction of neighbourhoods with node dynamics. The communicability centrality [Estrada et al., 2009] is a lesion based measure based on the notion of communicability [Estrada and Hatano, 2008, Andreotti et al., 2014], that particularly applies to broadcasting, and quantifies the global effect of node removal on network communication. Communicability and its associated centrality measure however, are static network quantities that do not take the dynamics into consideration. Flow on the other hand is a dynamics-dependent quantity that adds a dimension of complexity to the static network based measures such as communicability centrality. The calculation of flow is also carried out in a neurodynamical system, which implies a broadcasting communication framework, thus making flow a general measure that makes no distinction between paths and requires information only of the local neighbourhood. The utility of flow lies in its ability to identify nodes through which the maximum amount of information flows, making them ideal targets to study or intervene and modulate the spread of network perturbations. The functional lesion of a node ‘i’, such that its activity is frozen at steady state, makes it “blind” to a perturbation, and results in the termination of the signal transfer further onwards from it. The effect of this exercise is the removal of the contribution of ‘i’ on the responses of the rest of the network, upon perturbation. The calculation of flow using equation (6) condenses the overall effect of this functional lesion of ‘i’ over all possible perturbations in the network, and thus equivalently all possible information transfer scenarios. The lesion of high flow nodes thus have the greatest impact on information flow in an active network, for the given dynamics. This can have particular applications in designing interventions to limit the the impact of perturbations to healthy brain function that can be brought about by neuropathologies, which can eventually result in a “new normal” (the disease state), similar to the perturbed steady state in this study. Flow can also find its use in the study of regions facilitating seizure propagation in epilepsy, using whole-brain epileptor models [Proix et al., 2017, Jirsa et al., 2017, El Houssaini et al., 2020].

Our results show that flow is maximum for nodes of intermediate strength, but close to the periphery (Fig.4). Comparing these nodes to the strength distribution (Fig.2B) shows us that these regions are also in much greater abundance than the extreme peripheral or hub nodes. A previous study by [Gollo et al., 2015] on the macaque network classified nodes into rich, feeder, and periphery depending on their degree of separation from the rich club. The feeder nodes were the most abundant in the network, and bridged the core to the periphery and were also shown to have faster dynamics than the structural core. It is also well established that the core and peripheral nodes have distinct information processing functions [Gollo et al., 2015, Bassett et al., 2013] and constantly communicate with each other. The intermediate strength dominance of flow in the empirical best fit model also suggests a polysynaptic parallel core-periphery communication pathway, and the abundance of the nodes suggest that the nodes are likely to fall into the “feeder” category. This can be alternatively viewed as a bidirectional core-intermediate-periphery pathway, with the intermediate strength nodes playing a central role. The flow structure below and above the best fit value of the scaling parameter shows a hub dominated and periphery dominated flow structure respectively, both of which don’t support a parallel core-periphery communication structure, and instead attributes importance in communication to the hub and the periphery (periphery-core-periphery or core-periphery-core pathways). Furthermore, although we look at the temporally asymptotic behaviour of regions in this work, the intermediate dominance of flow would additionally imply that these regions display fast dynamics [Gollo et al., 2015], which we note would be necessary for efficient and fast communication between distant regions. The network level flow analysis shows that the regions that comprise the Salience/Ventral Attention network (SNVA) have the highest average flow values, implying that the SNVA plays the most important role in mediating information flow between arbitrary pairs of brain regions outside the network. Networks with high flow can be viewed as “gatekeeper” networks, capable of strongly regulating the information flow between other networks. This result additionally offers a network-dynamical take on Menon’s triple network model which implicates the SNVA as a major network mediating information flow between the Frontoparietal control network (Con) and the Default Mode Network (DMN), the failure of which results in major neuropathologies [Menon, 2011].

The observation that regions with high flow are also likely to show a net negative response asymmetry, i.e. response to external stimulations is greater than response elicited, is a fallout of flow’s dependence on the linear response matrix. The activity of regions with a high negative influence can be altered more by perturbations originating from the rest of the network, when compared to that of regions with lesser negative influence. This directly translates to a greater contribution by these negative influence regions to the response of a node to a perturbation that originated from an arbitrary node.

### Methodological considerations

Some points have to be considered regarding this formalism, in order to truly understand its scope, and work on possible limitations. *First*, while the formalism is general enough to be applied on any scale, the distribution of network influence and flow is entirely dynamics dependent. *Second*, in translating this technique into whole-brain perturbation-response experiments, executing functional lesions will require some innovation, in order to probe flow. One possible method suggested earlier [Harush and Barzel, 2017] is to carry out a negative perturbation on the node to be lesioned, such that its activity is maintained at the original baseline. *Third*, the analyses here are carried out on noiseless models, as we are interested in the overall change of amplitude (activation) of signals, which is sufficient for a proof of concept. The formalism can be readily applied to a stable stochastic system as well. *Fourth*, the perturbations are assumed to be small enough to elicit linear responses.

This work presents a simple scale-invariant perturbative formalism that can be used to both probe the distributions of influential nodes in the network, as well as identify nodes that are central to communication processes, both of which have immense experimental and clinical importance in neuroscience. The robust theoretical framework surrounding the formalism additionally opens up possibilities of mathematically studying the communication patterns permitted by established neurodynamical models. Most importantly, the formalism is invariant to the form of the underlying dynamics and instead gathers insights purely based on the response of a perturbed system of neuronal entities, making it ideal for implementation with minimal assumptions about how the system functions.

## Abbreviations

Con: Frontoparietal Control network
DA: Dorsal Attention network
DMN: Default Mode network
DTI: Diffusion Tensor Imaging
DWI: Diffusion Weighted Imaging
Lim: Limbic network
LSM: Linear Stochastic Model
MFM: Mean Field Model
rs-FC: Resting state Functional Connectivity
RSN: Resting State Network
SC: Structural Connectivity
SMot: Somato-motor network
SNVA: Salience/Ventral Attention network
Vis: Visual network

## Acknowledgments

We acknowledge the generous support of NBRC Core funds and the Computing facility. For simulations, resources from the Neuroscience Gateway [Sivagnanam et al., 2013] were used. Data collection and sharing for this project was provided by the Cambridge Centre for Ageing and Neuroscience (Cam-CAN). Cam-CAN was supported by the UK Biotechnology and Biological Sciences Research Council (Grant BB/H008217/1), together with support from the UK Medical Research Council and University of Cambridge, UK. In accordance with the data usage agreement for Cam-CAN dataset, the article has been submitted as open access. The preprocessed Nathan Kline Institute (NKI) Rockland sample dataset was obtained from the publicly accessible UCLA Multimodal Connectivity Database (UMCD) [Brown et al., 2012]. The NKI data can also be obtained from the International Neuroimaging Data-Sharing Initiative (INDI) website (http://fcon_1000.projects.nitrc.org/indi/pro/nki.html).

## Funding information

Arpan Banerjee, Ministry of Youth Affairs and Sports, Government of India, Award ID: F.NO.K- 15015/42/2018/SP-V. Arpan Banerjee, NBRC Flagship program, Department of Biotechnology, Government of India, Award ID: BT/MED-III/NBRC/Flagship/Flagship2019.

## Footnotes

1 For this study, “network” refers to a physically connected network

2 In an experimental setting, this response matrix would be populated by the observed responses of regions to perturbations, as given by (2)

## Notes

### Competing Interest Statement

The authors have declared no competing interest.

