## Supplementary figures for "A scale-invariant perturbative approach to study information communication in dynamic brain networks"

Fig. S1

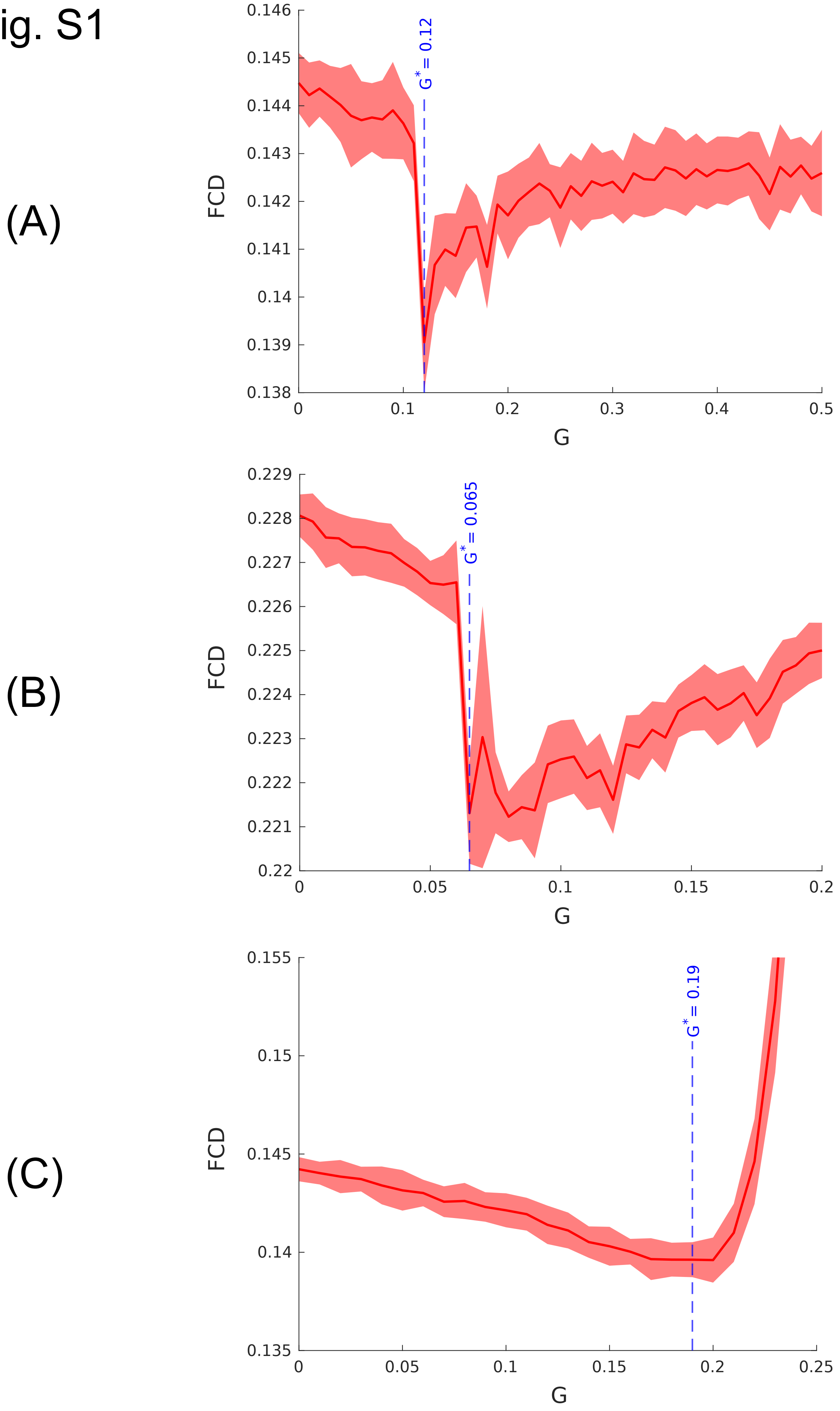


**Figure S1. Identification of model parameters that maximize similarity to empirical observations**

Identification of the model which best matched with the empirical ground-truth is depicted as a plot of the Functional Connectivity Distance (FCD) as a function of the free model paramter *G* that scales the structural connectivity (SC). The parameter value which corresponds to the best fit model minimises the FCD between the model FC and the empirical FC, and is denoted by the dashed line. The figures display the critical parameter values for **(A)** Mean Field Model (MFM) evolving on the Cam-CAN SC; **(B)** MFM evolving on the NKI SC; and **(C)** Linear Stochastic Model (LSM) on the Cam-CAN SC.

Fig. S2

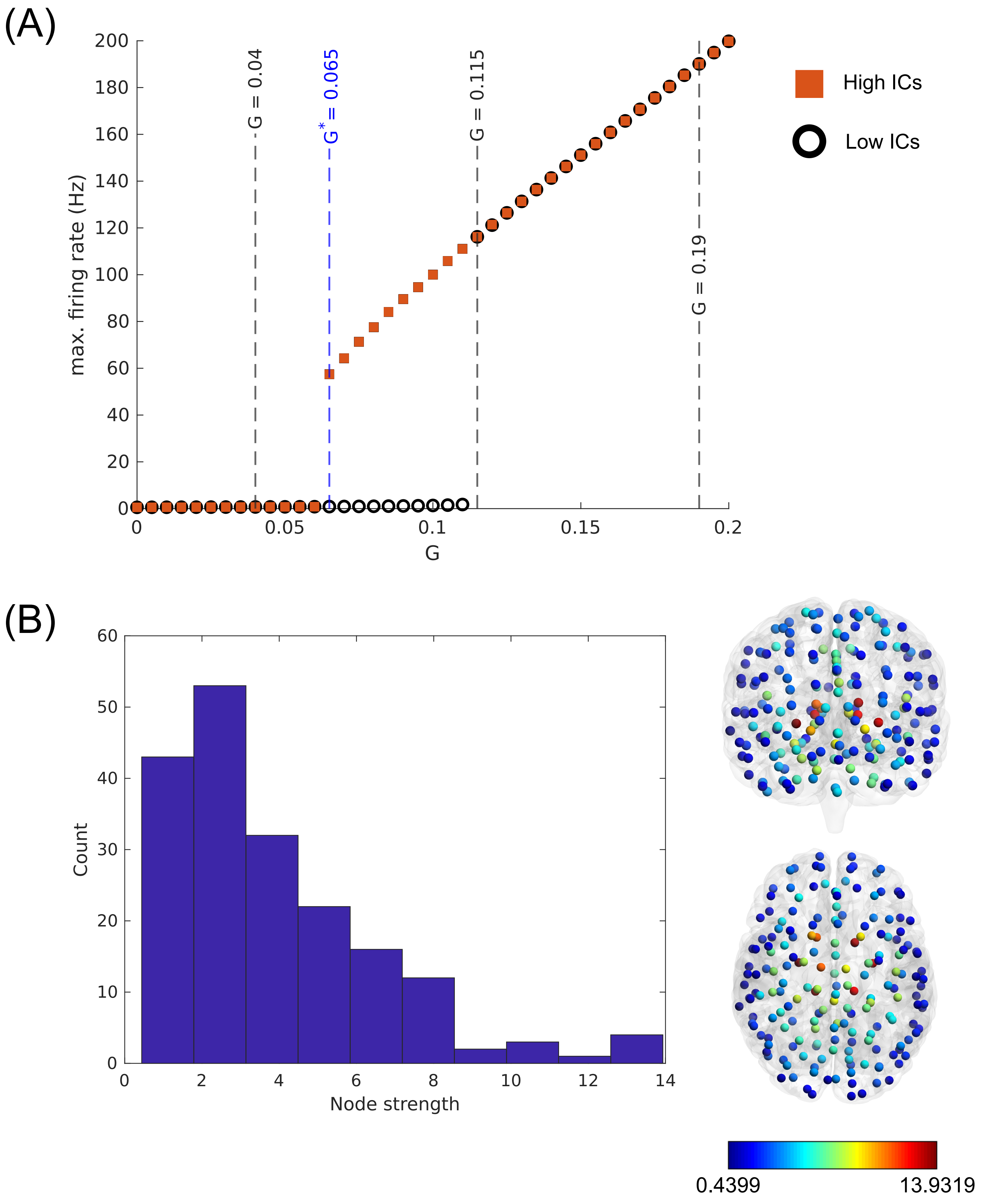


**Figure S2. MFM dynamical landscape for the NKI dataset**

**(A)** The figure shows the dynamical landscape of the MFM as a function of the global scaling parameter *G* (orange squares = High initial conditions(ICs) (0.3 ≤ S(t=0) ≤ 1.0), black rings = low ICs(0 ≤ S(t=0) ≤ 0.1)), quantified as the maximum firing rate among all the nodes in the network. Figure shows results for 10 random trials at each value of *G* ($\Delta G$ = 0.005). The values of *G* for which the response asymmetries and flows are calculated are marked by the dashed lines. The blue dashed line corresponding to $G^*$ = 0.065 is the value of $G$ for which the model maximally conforms to empirical data. **(B)** Node strength (sum of the number of white matter tracts between the node and its neighbours) distribution of the NKI dataset (*left*) with 188 nodes of the Craddock-200 atlas, and associated BrainNet plots where colour maps to node strength (*right*).

Fig.S3

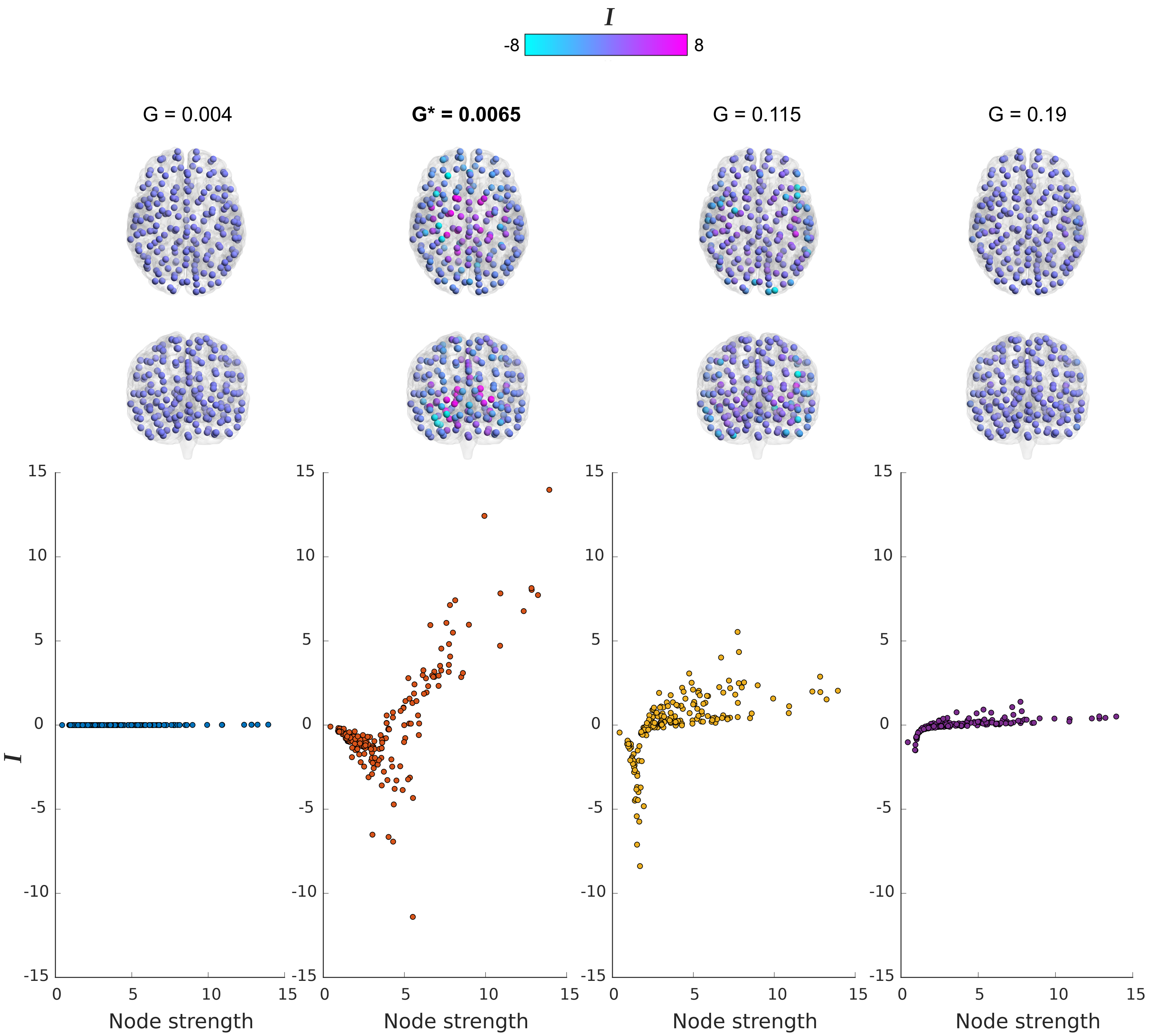


**Figure S3. Response asymmetry in the NKI dataset**

Response asymmetries, quantified as the net influence of nodes, *I*, as a function of node strengths, for nodes belonging to the NKI dataset. The values of the global scaling parameter for which the asymmetries are computed are shown in Fig.S2. Anterior and ventral view BrainNet plots depict the inter-hemispheric symmetry of net influence distribution, and the colour maps to the associated value (maximum colour value set to 8 for clarity). Response asymmetry is maximized at $G^* = 0.065$ and reduces as $G$ is increased.

### Fig. S4

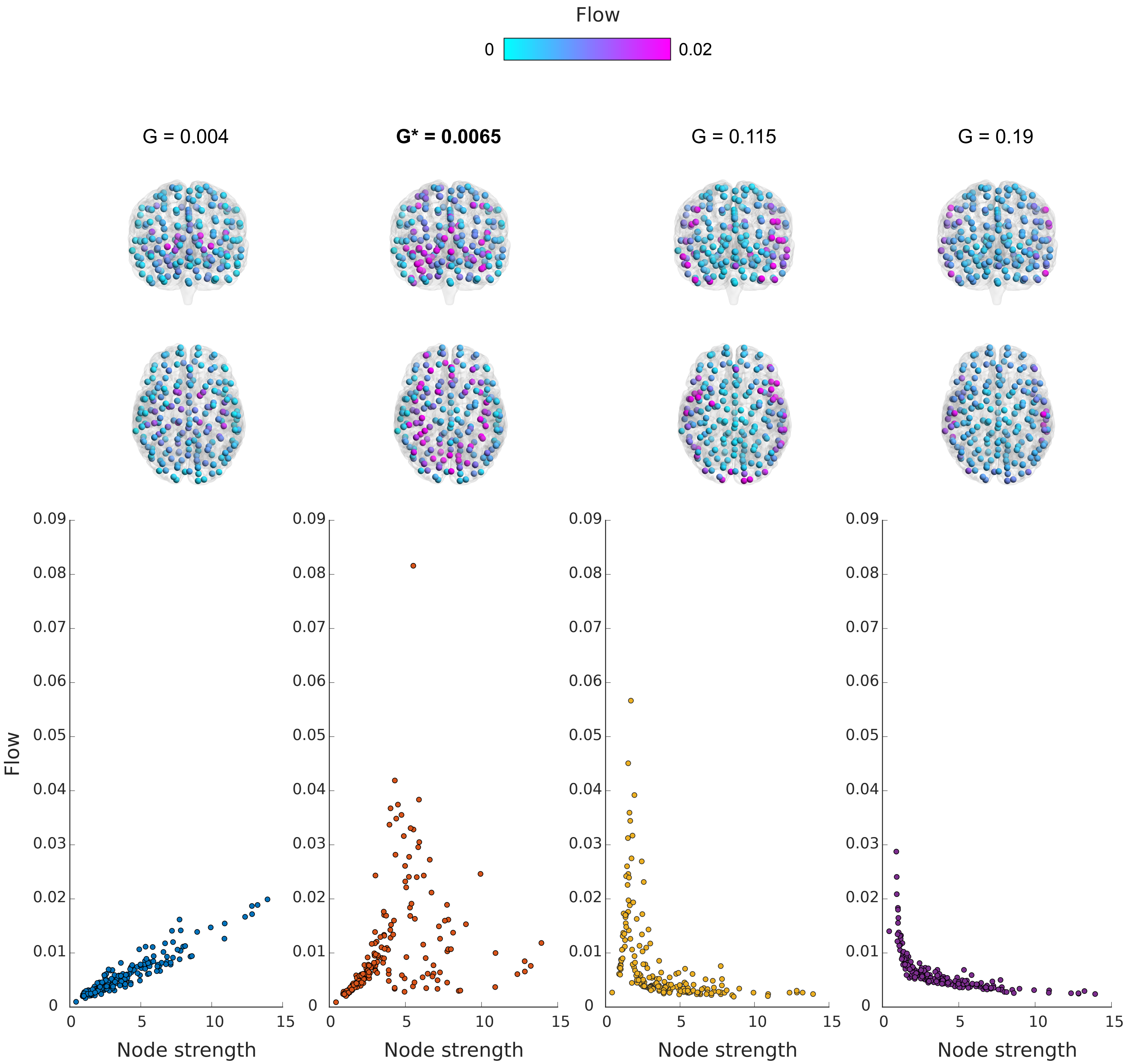


**Figure S4. Node-level flow structure in the NKI dataset**

Flow through a node, quantified as the effect of freezing the activity of node on the magnitude of responses elicited on the rest of the network, as a function of node strengths, for nodes belonging to the NKI dataset. The values of the global scaling parameter for which the flow values are computed are shown in Fig.S2. Anterior and ventral view BrainNet plots depict the inter-hemispheric symmetry of flow distribution, and the colour maps to the associated value (maximum colour value set to 0.02 for clarity). Flow is maximized at intermediate strength nodes for $G^{*} = 0.065$, but moves further towards the periphery (low strength nodes) as $G$ is increased.

### Fig. S5

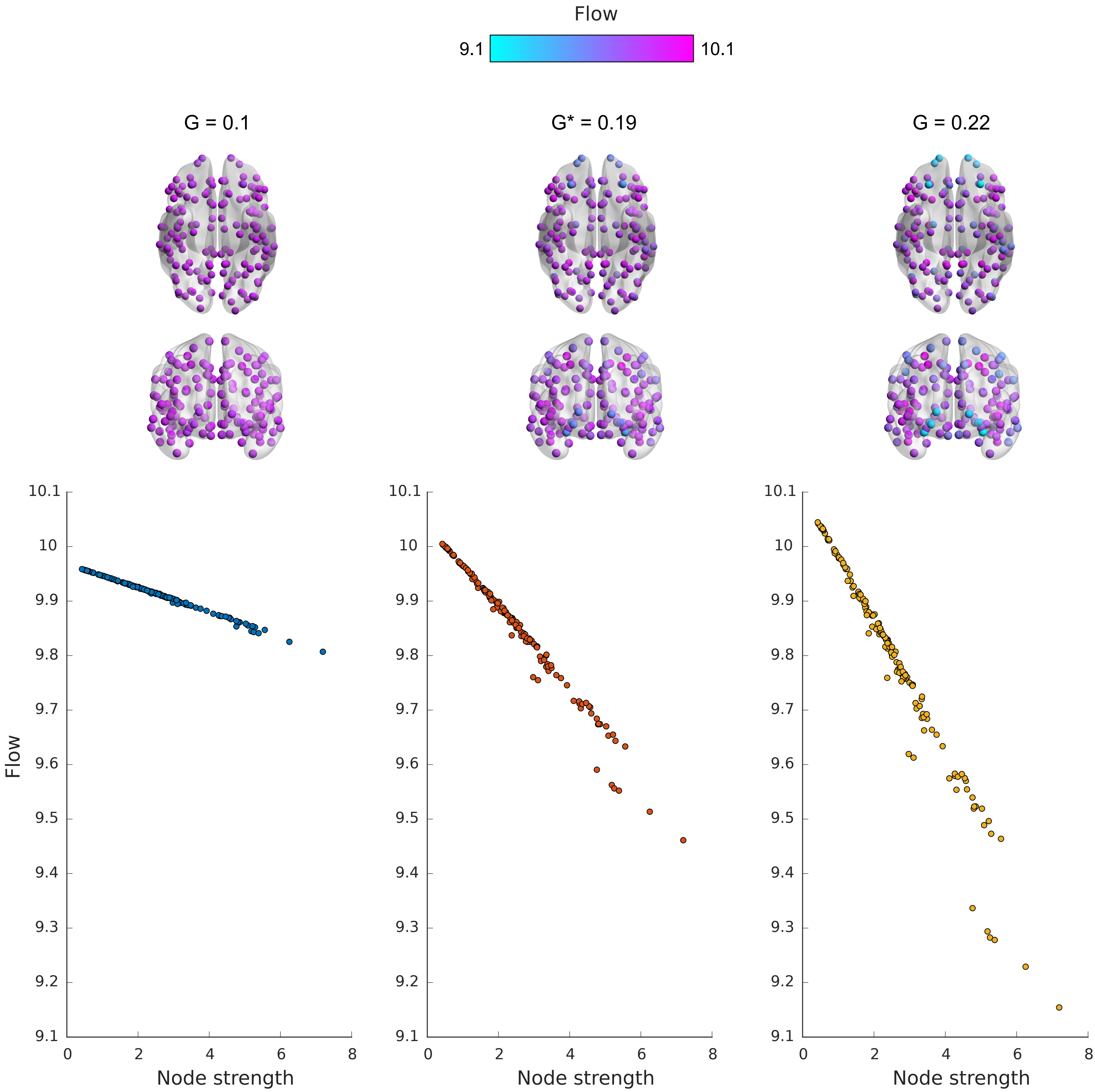


**Figure S5. Node-level flow structure for the LSM on the Cam-CAN dataset**

Flow through a node as a function of node strengths, for nodes following LSM dynamics, evolving over the SC derived from the Cam-CAN dataset. The values of the global scaling parameter for which the flow values are computed are centred around the empirical best fit for the LSM, $G^{*}$=0.19 , as shown in Fig.S1C. Anterior and ventral view BrainNet plots depict the inter-hemispheric symmetry of flow distribution, and the colour maps to the associated value of flow. The flow is clearly periphery dominated for all the values of $G$, and the range of the distribution of flow values increases with $G$.
